# Molecular and cell phenotype programs in oral epithelial cells directed by co-exposure to arsenic and smokeless tobacco

**DOI:** 10.1101/2024.10.14.618077

**Authors:** Samrat Das, Shefali Thakur, Vincent Cahais, François Virard, Liesel Claeys, Claire Renard, Cyrille Cuenin, Marie-Pierre Cros, Stéphane Keïta, Assunta Venuti, Cécilia Sirand, Akram Ghantous, Zdenko Herceg, Michael Korenjak, Jiri Zavadil

**Affiliations:** Epigenomics and Mechanisms Branch, International Agency for Research on Cancer, Lyon, France; The Breast Cancer Now Toby Robins Research Centre, The Institute of Cancer Research, London, UK; University Claude Bernard Lyon 1, INSERM U1052–CNRS UMR5286, Cancer Research Center, Centre Léon Bérard, Lyon, France; University of Lyon, Faculty of Odontology, Hospices Civils de Lyon, Lyon, France; Centre of Excellence in Mycotoxicology and Public Health, Faculty of Pharmaceutical Sciences, Ghent University, Ghent, Belgium

**Keywords:** arsenic, smokeless tobacco, oral cells, DNA methylation, gene expression, whole-exome sequencing, live cell analysis, apoptosis, inflammatory response

## Abstract

Chronic arsenic exposure can lead to various health issues, including cancer. Concerns have been mounting about the enhancement of arsenic toxicity through co-exposure to various prevalent lifestyle habits. Smokeless tobacco products are commonly consumed in South Asian countries, where their use frequently co-occurs with exposure to arsenic from contaminated groundwater. To decipher the *in vitro* molecular and cellular responses to arsenic and/or smokeless tobacco, we performed temporal multi-omics analysis of the transcriptome and DNA methylome remodelling in exposed hTERT-immortalized human normal oral keratinocytes (NOK), as well as arsenic and/or smokeless tobacco genotoxicity and mutagenicity investigations in NOK cells and in human p53 knock-in murine embryonic fibroblasts (Hupki MEF). RNAseq results from acute exposures to arsenic alone and in combination with smokeless tobacco extract revealed upregulation of genes with roles in cell cycle changes, apoptosis and inflammation responses. This was in keeping with global DNA hypomethylation affecting genes involved in the same processes in response to chronic treatment in NOK cells. At the phenotypic level, we observed a dose-dependent decrease in NOK cell viability, induction of DNA damage, cell cycle changes and increased apoptosis, with the most pronounced effects observed under arsenic and SLT co-exposure conditions. Live-cell imaging experiments indicated that the DNA damage likely resulted from induction of apoptosis, an observation validated by a lack of exome-wide mutagenesis in response to chronic exposure to arsenic and/or smokeless tobacco. In sum, our integrative omics study provides novel insights into the acute and chronic responses to arsenic and smokeless tobacco (co-)exposure, with both types of responses converging on several key mechanisms associated with cancer hallmark processes. The generated rich catalogue of molecular programs in oral cells regulated by arsenic and smokeless tobacco (co-)exposure may provide bases for future development of biomarkers for use in molecular epidemiology studies of exposed populations at risk of developing oral cancer.

## 1. INTRODUCTION

Arsenic contamination of groundwater is a major public health problem globally (1). Long-term consumption of arsenic via drinking water has the potential to cause serious adverse health effects including cancer (2–8). Moreover, there has been a growing concern about co-exposure to various prevalent lifestyle factors and their role in the enhancement of arsenic toxicity(9, 10). Habitual consumption of smokeless tobacco in many Asian countries is one of such factors. SLT is thought to be a major risk factor for the high incidence of head and neck cancers (11–16). Smokeless tobacco products are commonly consumed in certain regions with confirmed arsenic contamination of the groundwater, such as in North-Eastern India (17, 18). *Sadagura* is a form of smokeless tobacco preparation that is widely consumed by the population in the southern part of Assam, India (19, 18).

The various biological processes associated with the initiation of oncogenesis due to arsenic and smokeless tobacco co-exposure remain to be fully understood. Genetic and epigenetic remodelling in cells induced by exogenous exposures to carcinogens contributes to carcinogenesis (20–23). Among the various epigenetic mechanisms underlying the cancer development, DNA methylation plays an important role, via the regulation of gene expression, with major implications for the initiation of the oncogenic process (24–35). Exposure to arsenic has been reported to cause changes in DNA methylation (36–41) and gene expression (42). Tobacco consumption-associated cancers have been mostly marked by genetic changes (43–48), although DNA methylation changes have also been implicated (49).

It has been suggested that tobacco consumption may aggravate arsenic-mediated toxicity (9). Studies in exposed animals (50, 51, 18, 52) and evaluation of samples from exposed human populations (17, 18) have indicated the DNA damage-causing potential of arsenic and smokeless tobacco co-exposure. However, global molecular alterations in the context of arsenic and smokeless tobacco co-exposure in relevant, experimentally controlled models have not been systematically characterized and there remains a need to better understand the various underlying biological processes associated with the initiation of oncogenesis.

In the current study we devised and employed a comprehensive multi-omics strategy consisting of analyses of the DNA methylome and transcriptome remodelling and of acquired genome-wide mutation spectra, combined with investigations of live-cell phenotypic readouts in appropriate *in-vitro* cell models. Using this integrative approach, we aimed to decipher both acute and long-term biological events and processes associated with arsenic and smokeless tobacco (co-)exposure in oral cells, and to understand the underlying molecular programs with potential relevance to oral cancer formation.

## 2. EXPERIMENTAL PROCEDURES, MATERIAL AND METHODS

### 2.1 Cells and cell culture

#### 2.1.1 hTERT-immortalized human Normal Oral Keratinocytes (NOK)

NOK cells were a kind gift from Dr Paul Lambert (University of Wisconsin-Madison, United States of America) (53, 54). The cells were propagated in the Keratinocyte growth medium 2 (PromoCell GmbH, Heidelberg, Germany) and 1% Penicillin/Streptomycin.

#### 2.1.2 Human p53 Knock-in Murine Embryonic Fibroblasts (Hupki MEF)

Primary Hupki MEF cells were received from the Laboratory of Dr Monica Hollstein (Department of Genetic Alterations in Carcinogenesis, German Cancer Research Center, Heidelberg, Germany) (55–59). Cells were maintained in complete medium consisting of Advanced DMEM (Gibco), 15% Foetal calf serum, 1% L-glutamine, 1% Penicillin/Streptomycin, 2% Pyruvate. Cells were passaged at nearly 70-80% confluency using 0.05% trypsin.

### 2.2 Test substances

Sodium arsenite (SA) solution (NaAsO₂, concentration of 0.05 mol/l (0.1 N), Cat. No. 106277) was purchased from Merck, Germany. From the stock solution of 0.05 mol/l, Intermediate stock solutions of 100 µM were prepared. From intermediate stock solution, the various final working dosage solutions were prepared by dissolving in appropriate culture medium volume. Smokeless tobacco aqueous extract was prepared as follows: The aqueous extract of *sadagura* smokeless tobacco preparation (SLT) was prepared following the method described by Jeng and co-workers (60) and with modifications (50, 51, 18). Briefly, all the ingredients namely, tobacco leaves, aniseeds, and black cumin seeds, were purchased from a local market in Silchar, Assam, India. Ingredients were independently dried, simmered and pounded in a blender to a fine powder. Afterwards, the ingredients were mixed in a suitable proportion based on weight (2:4:1 ratio) and dissolved in milliQ water in a shaker with continuous stirring for 72 h (at 4°C). Afterwards, using clean cheesecloth, debris was removed, and the resulting filtrate was subsequently passed through Whatman filter paper No. 1. The extract was dried and was subsequently preserved in a freezer (–20°C). Fresh aliquots of specific doses/concentrations were prepared prior to the *in vitro* treatment by dissolving appropriate amounts of extract in milliQ water subsequently passed through 0.20 µM syringe filters (Sartorius, Germany). The method for testing of various concentrations and for the current study have been presented in section 2.6. Half maximal inhibitory concentrations (IC_50_) obtained were used for determining appropriate co-exposure dose(s) for analyses of the transcriptome (NOK cells), DNA methylome (NOK cells), γH2AX immunofluorescence (NOK and Hupki MEF cells) and genome-wide mutagenesis (NOK and Hupki MEF cells).

To investigate the possible potential synergistic co-exposure effects, exact additive dosage composed of SA and SLT individual doses were selected for the comet assay, apoptosis analysis, cell cycle analysis and Incucyte live cell monitoring studies.

### 2.3 Temporal RNA-seq analysis of exposed NOK cells

NOK cells were grown in T25 flasks (seeding density ∼ 3 - 5 × 10^5^ cells). Upon reaching a∼ 70% confluency the cells were treated with media containing the test substances with doses as indicated in corresponding figure captions. The co-exposure doses were selected taking into consideration of the ratio of IC_50_ of the individual exposures. The experimental groups consisted of various time points based on exposure period (2, 4 and 8 h). Two experimental control groups were included, the baseline untreated control (0 h), and time-matched untreated control (harvested at 8 h). Each timepoint condition consisted of four experimental replicates. After exposure, the media containing the test substances were removed from the flasks and the cellular RNA was extracted using TRIzol^TM^ Reagent (Cat. No. 15596026, Invitrogen). Estimation of yield was done by Qubit analysis (Qubit 3.0 Fluorometer, Life Technologies, Invitrogen™, Thermo Scientific Fisher). RNA integrity was analysed on High Sensitivity RNA ScreenTape (Cat. No. 5067-5579, Agilent), using the 4200 TapeStation automated electrophoresis instrument (Agilent). All samples exhibited RNA Integrity Number (RIN) of 8.4 or above.

500 ng RNA per sample was used to prepare libraries (QuantSeq 3’ mRNA-Seq Library Prep Kit for Illumina (FWD)), Cat. No. 015, Lexogen). Multiplexed samples were sequenced as 75 bp single reads on the Illumina NextSeq 500 platform. Reads were trimmed and aligned against the human genome (build hg38). The data was analysed using the RNAseq-nf pipeline (https://github.com/IARCbioinfo/RNAseq-nf), and STAR (2.7.3.a), htseq (0.12.4), and DESeq2 (3.14) tools were employed for analyses (61). Quantile-normalized counts were pre-filtered (genes with read counts <200 were removed), log2 transformed, and the resulting data used for differential gene expression analysis. The gradual gene expression changes observed under the various experimental conditions were investigated using the Pavlidis Template Matching (PTM) analysis tool followed by hierarchical clustering of the obtained expression profile groups, using the TMeV suite (p<0.05) (62, 63).

### 2.4 DNA methylome analyses in NOK cells

The NOK cells underwent controlled treatment as follows: each week the cells were exposed for 24 h with doses as indicated in corresponding figure captions. The SA+SLT co-exposure doses were selected taking into consideration of the ratio of IC_50_ of the individual exposures. After the exposure period, the exposure medium was replaced with normal media and cells were allowed to recover. Exposures were repeated for a period of 5 weeks. Following the repeated exposure protocol, genomic DNA was extracted from the cells as described in section 2.5.

500 ng of DNA per sample were bisulfite-converted to enable quantitative assessment of the methylation levels over 850K CpG sites. The methylation profiles were obtained using the Infinium® Methylation EPIC bead arrays (Illumina), with recommended protocols applied for amplification, labelling, hybridisation, and scanning. Experimental triplicates were analysed for each treatment condition.

The raw data were processed using the *methylkey* pipeline (https://github.com/IARCbioinfo/methylkey) and *minfi* tool (3.14) was used (64). ProbeIDs were filtered, followed by overlap analyses of significantly methylated probes and the corresponding DMRs. Information of the closest genes and the associated delta beta values, chromosome location information (DMRs) with respect to a probe was clubbed with Illumina manifest file. The hypo-/hyper-methylated CpGs screened under various treatment specific conditions have been presented as heat maps. TMeV suite was used for heatmap figures (62). To study the effect of methylation on gene regulation in detail we focused on CpGs associated with promoter regions (1,500 bp upstream of the TSS). For biological processes/pathway analysis, NIH-DAVID tool was used and only the genes corresponding to the UCSC database column of the above-mentioned manifest file were considered (65).

### 2.5 Extraction, quantification, and quality control of the genomic DNA from cells

Genomic DNA from the cells was extracted using the NucleoSpin Tissue kit (Macherey-Nagel). The extracted DNA was then quantified by using Qubit dsDNA HS Assay Kits (Life Technologies, Invitrogen™, Thermo Fisher Scientific) in a Qubit 3.0 Fluorometer (Life Technologies, Invitrogen™, Thermo Fisher Scientific). The purity of the DNA samples were analysed by using ND-8000 spectrophotometer (NanoDrop Technologies Inc.). Further the DNA samples were also analysed by the 4200 TapeStation automated electrophoresis instrument (Agilent) for quality/fragment size assessment.

### 2.6 MTS cell viability assay and selection of exposure dosages

MTS assay was used to measure the metabolic activity as a proxy for cell viability upon exposure to increasing concentrations of the test compounds in NOK and Hupki MEF cells. This step helped determine the concentrations for the acute (NOK transcriptomics, Hupki MEF exome) and repeated (NOK DNA methylome & NOK exome) exposure experiments. The cells (5,000 cells/well) were seeded in 96 well plates and were subsequently treated with selected doses as indicated. The co-exposure doses were selected based on the ratio of IC_50_s of the individual exposures. Separate plates were established for 24 and 48 h time points. After the treatment period the cell viability was measured using the CellTiter 96 AQueous One Solution Cell Proliferation Assay (Cat. No. G3582, Promega Corporation). Briefly, the plates were incubated (3 h, 37°C) following which absorbance was measured at 492 nm (Apollo 11 LB913 plate reader). For each experimental condition the assay was performed in triplicates.

### 2.7 Phospho-H2AFX immunofluorescence in NOK and Hupki MEF cells

Immunofluorescence staining of phosphorylated histone H2AFX (γH2AFX) was performed using P-Histone H2A.X (S139) (20E3) Rabbit mAb (Cat No. 9718S, Cell Signaling Technology, Inc.). Briefly, cells were seeded in duplicates on coverslips in two different 24-well plates (for 24h and 48h conditions respectively) and, the following day, treated with optimized doses (indicated in the corresponding figure captions). The co-exposure doses were selected based on the ratio of IC_50_ of the individual exposures. Following the exposure period, the treatment medium was replaced with fresh medium in the wells and was kept for 4 h for recovery. Afterwards, the cells were fixed (4% formaldehyde, 20 min at room temperature). Following blocking in image-iT^TM^ FX signal enhancer (Cat No. A31627 B, Invitrogen by Thermo fisher scientific, Life Technologies corporation) for 60 min, they were incubated with the γH2AFX-antibody (1:500 in PBS) overnight at 4°C. Subsequent incubation with a fluorochrome-conjugated secondary antibody Alexa Fluor^TM^ 488 goat anti-rabbit (Cat No. A31627 A, Invitrogen by Thermo fisher scientific, Life Technologies corporation) was performed for 60 min at room temperature. Coverslips were mounted in Vectashield antifade mounting medium with DAPI (Ref: H-1200, Vector Laboratories). The coverslips were analysed, and images were captured in fluorescent microscope (Nikon Eclipse Ti).

### 2.8 Comet assay in NOK cells

Comet assay was performed to determine the extent of DNA damage in acutely exposed cells to varying doses (refer to the corresponding figure captions). Additive dosage was selected for co-exposure to investigate the potential synergistic effects. Briefly, 5-10 µL of the sample was taken and properly mixed with 70-75 mL LMPA (Low melting point agarose) and rapidly spread on precoated (1% normal melting agarose) frosted slides. Slides were kept in the ice pack for 5–10 min, afterward, a third agarose layer of 80 µL LMPA was added to the slides. Slides were then immersed in cold lysis solution (pH 10) and kept overnight at 4°C. After lysis step, the slides were kept in the electrophoresis buffer (300 mM NaOH: 1 mM Na_2_EDTA at pH 13.5) for 20 min at 4°C to allow unwinding of the DNA. Following unwinding, electrophoresis of the slides was conducted (24 V, 300 mA, 4°C, 30 min). Slides were neutralized in 0.4 M Tris buffer (pH 7.5) 3 times for 5 min, followed by staining with ethidium bromide. Comet slides images were captured using fluorescence microscope (Nikon Eclipse Ti). Images of 100 cells in triplicate per condition were analysed using “OpenComet version 1.3.1” software tool (66).

### 2.9 Generation of clonal population in exposed NOK and Hupki MEF cells for whole-exome sequencing

#### 2.9.1. Single cell subcloning of NOK cells

For single cell subcloning of NOK cells (pre- and post-treatment), briefly, flasks containing cells were trypsinized at 70-80% confluency. Next the cell solution was passed through a cell strainer to eliminate cell clumps and break up clusters of cells into single cells. The concentration of viable cells in the filtered solution was assessed using a cell counter (TC-20, Bio-rad) and trypan blue staining. The cells were diluted in multiple steps to set up a final seeding requirement of 0.5 cells/well in 96-well plates.

#### 2.9.2 Barrier bypass-clonal expansion of Hupki MEF cells

Barrier bypass-clonal expansion of Hupki MEF cells was done following the established protocols (55, 56, 67, 59). Briefly, both untreated as well as treated primary cells were cultured until they bypassed the senescence stage and immortalized clonal cell populations emerged. The primary cells used as baseline controls and the resulting clones were compared in WES analysis (see below).

#### 2.10 Whole-exome sequencing of exposed NOK and Hupki MEF cells

DNA library preparation was carried out using the KAPA HyperPlus kit (KAPA Biosystems) according to the manufacturer’s instructions. Exome capture was performed using the SeqCap EZ Mouse or Human Exome v3 kit (Roche). Briefly, an equimolar pool of DNA samples (total 1 µg) was prepared. To this pool, 5 µL of COT (human or mouse, depending on the sample type. COT is used to block non-specific hybridization. Human COT DNA was from Roche (accessory kit), and the Mouse COT DNA was from Life Technologies), 1 µl of IDT blocking oligo i7 and 1 µl of IDT blocking oligo i5 (used to bind to adapter sequences to help prevent non-specific binding) was added. The required volume of the AMPure XP (Beckman Coulter) beads were added, and incubated up to 15 min at room temperature, and then washed twice with 80% EtOH and eluted in a mix of 7.5 µl hybridization buffer and 3 µl Component A (Roche accessory kit). Followed by further incubation at 95 °C for 10 min, and transfer of the pool in a tube of capture probes. Incubation of 72h at 47 °C was performed and later bead capture and washing steps were followed by using the Roche SeqCap EZ Hybridization & Wash kit (Roche). Exome-captured libraries were sequenced on the NextSeq 500 platform (Illumina, Inc.) in a 75 bp paired-end read mode.

To detect single base substitution (SBS) mutations from the data generated after chronic exposure, two somatic variant callers were used, namely MuTect2 (68) and Strelka2 (69). The vcf files were annotated using the gama_annot-nf pipeline (https://github.com/IARCbioinfo/gama_annot-nf) and after a pre-processing step that included the merging of all sample files for both callers (independently), the data was filtered to remove variants with low coverage, mutations in repeated regions or single nucleotide polymorphisms (SNPs). The union of the mutations passing filters for both caller was retained.

#### 2.11 Cell cycle analysis in NOK cells

Following acute exposure (24h/48h) of the test substances in the experimental cells (doses indicated in corresponding figure captions, with the additive co-exposure dosage selected to address potential synergistic effect) , cell cycle analyses was done by using propidium iodide staining and analysing the samples in flow cytometer (BD FACSCanto^TM^ II). Briefly, the cells were washed in PBS and centrifuged at 1600 rpm for 5 min to collect the cell pellets. 1mL of cold 70 % ethanol was added, and the cells were fixed, vortexed and incubated in ice for 30 min. The cells were pelleted and washed with PBS 3 times. To each sample, 500 µL of PBS +PI (25 µg/ ml) + RNAase (100 µg/ ml) was added and mixed properly, followed by incubation (30 min, in a rotating wheel in dark). Then cells were further incubated at 4°C for 90 min. Following which the samples were analysed in the flow-cytometer. 10,000 events were analysed in triplicates per experimental treatment condition.

#### 2.12 Apoptosis study in NOK cells

Following acute exposure of cells to test substances (24h/48h, with doses indicated in corresponding figures), apoptosis study was performed by using PE Annexin V apoptosis detection Kit I (Cat. 559763, BD Pharmingen^TM^) and analysing the samples in flow cytometer (BD FACSCanto^TM^ II). For performing the assay, the protocol mentioned in the kit was followed. Briefly the cells were washed with cold PBS and re-suspended in 1X buffer (1×10^6^ cells/mL). 100 µL of the solution (1 × 10^5^ cells) were transferred to a 5 mL tube, to which 5 µL of PE Annexin V and 5µL of 7-AAD were added. The tube was gently vortex and incubated for 15 min (room temperature, dark conditions). Subsequently, 400 µL of 1X binding buffer was added to each tube and the samples were analysed by flow cytometer immediately. 10,000 events were analysed in triplicates per experimental treatment condition.

#### 2.13 Monitoring the live cell death and confluency in NOK cells

24-well plates were employed and 0.05×10^6^ cells were seeded each well and kept in the cell culture incubator for two days. After two days, respective treatments, with doses indicated in corresponding figure captions, were administered in the wells and the plate was immediately transferred to Incucyte S3 live-cell analysis instrument (Essen BioSciences, Sartorius) for live monitoring of the cell growth till 48h. Each treatment condition was supplemented with or without Caspase Inhibitor Z-VAD-FMK (Cat. No. FMK001, R&D Systems, Inc.). Each experimental condition group had wells in duplicate. Live monitoring data was processed by IncuCyte® Analysis Software (Essen BioSciences, Sartorius).

## 3. RESULTS

### 3.1. Transcriptomic responses to arsenic and SLT exposure in NOK cells

To understand how arsenic and SLT exposure(s) affect the gene expression responses, we performed RNA-seq analysis on normal oral keratinocytes (NOK) exposed for up to 8 hours. The treatment-specific modulation of gene expression is shown in Figure 1 and the associated, selected biological processes are summarized in Supplementary Table 1. Gene expression was reproducibly affected by SA (188 genes associated with SA-exposure specific upregulation; 34 genes associated with SA-exposure specific downregulation) and SLT treatment (123 genes associated with SLT-exposure specific upregulation).

**Figure 1.**
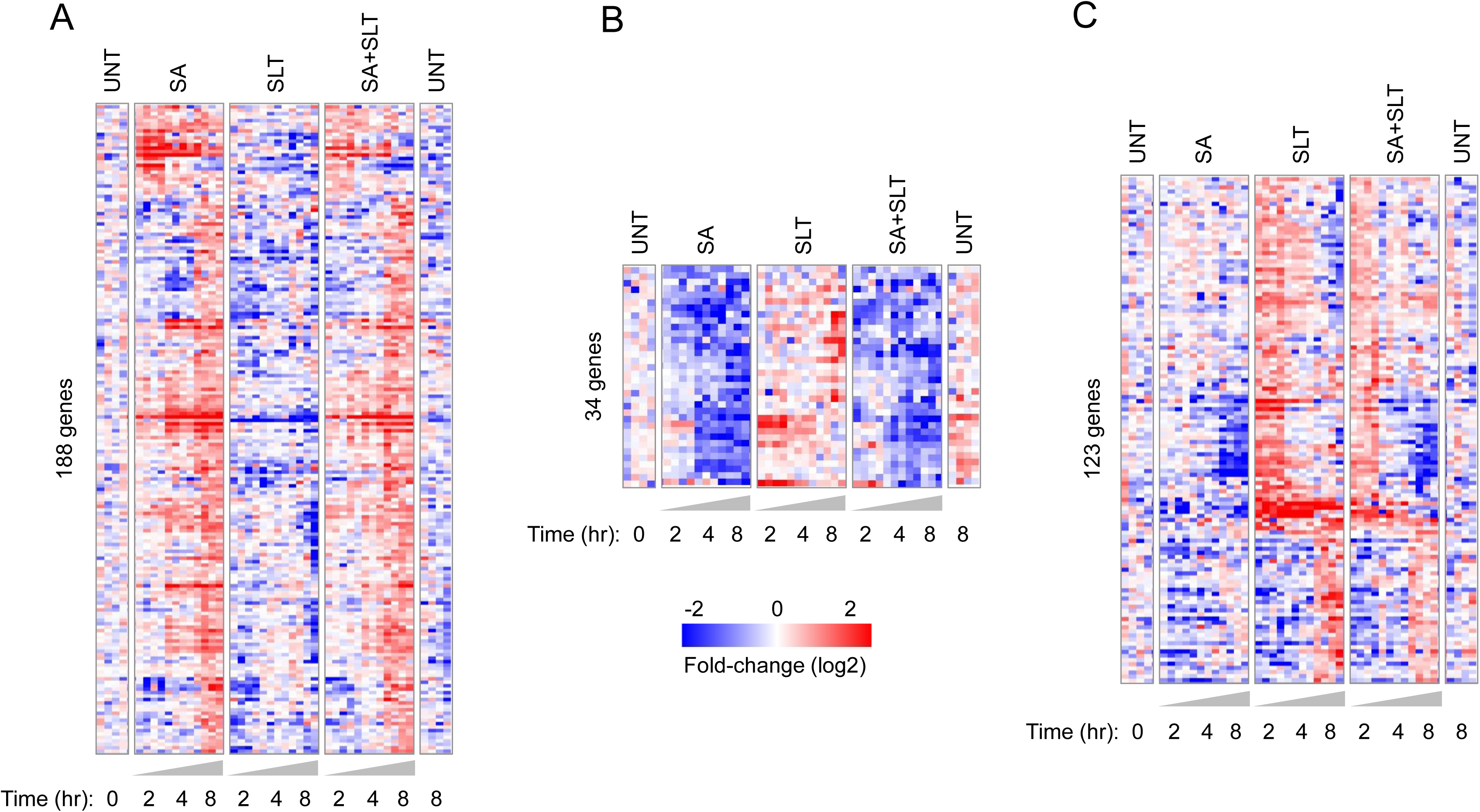
Gene regulation patterns specific to treatment conditions in NOK cells (at 2, 4, 8h). A) SA Upregulated genes; B) SA downregulated genes; C) SLT Upregulated genes. Abbreviations: UNT – Untreated time matched controls, SA - Sodium arsenite (10 µM), SLT - Smokeless tobacco extract (3000 µg/mL), SA+SLT - Sodium arsenite (6.5 µM) + Smokeless tobacco extract (750 µg/mL).

Around 30% (55 out of 188) genes were upregulated by SA in an immediate to intermediate early response manner, a pattern enriched for genes active in metabolic pathways (*ATP6VIC1, CERS3, GBA3, PI4K2A, PNP, SMOX*). Other crucial biological processes associated with SA-specific gene upregulation were cell cycle regulation, response to oxidative stress and DNA damage and inflammatory response. (Fig. 1, Supplementary Table 1a). Several biological pathways associated primarily with SA-specific downregulation consisted of regulation of transcription from RNA polymerase II promoter, wound healing, WNT signaling pathway, proteolysis, regulation of transcription (DNA-templated), and others (Supplementary Table 1). Notably, we observed other affected programs of systemic relevance, including pathways in cancer, cellular senescence, cell cycle checkpoints, cell cycle, immune system and others. (Fig. 1, Supplementary Table 1b). Around 67% (82 out of 123) genes were responding to SLT alone in an immediate to intermediate fashion, with some of these genes playing roles in important pathways such as p53 signalling pathway (*ATM, TNFRSF10B, CCNG1, SESN3),* and FoxO signaling pathway *(ATM, FOXO4, CDKN2B, IRS1*), microRNAs in cancer (*ABL1, ATM, CCNG1, IRS1*).

Among the notable biological processes associated with SLT-specific upregulation were regulation of cell cycle, cellular response to DNA damage stimulus, inflammatory response, apoptotic process, cellular response to oxidative stress and chromatin remodelling (Supplementary Table 1). The prominent SLT-modulated pathways were related to immune system, pathways in cancer, chemical carcinogenesis - reactive oxygen species, apoptosis, DNA double-strand break repair, and cell cycle checkpoints (Fig.1, Supplementary Table 1c). The general background gene regulation patterns shared across all treatment conditions (i.e. irrespective of exposure type) are presented in Supplementary Figure 1.

Our results show that SA and SLT exposure induce marked exposure specific as well as non-specific changes in gene expression, indicating that exposure to SA and/or SLT affects important cancer-related processes including the regulation of cell cycle, cell death and inflammatory responses.

### 3.2 Arsenic and SLT induce DNA methylome remodelling in NOK cells

Findings from the transcriptomic analysis prompted us to investigate whether SA and SLT-induced epigenetic changes would affect similar or different biological processes. We performed chronic exposure experiments in NOK cells and carried out global DNA methylome analysis (Fig. 2A). Overall, SA and SA+SLT treatment resulted in more hypo-than hypermethylation (Fig. 2B-e, Fig. 2B-d). The observation of 3,948 hypomethylation events upon SA+SLT co-exposure not detected in the single treatment conditions indicated specific DNA methylome remodelling solely due to the combined exposure (Fig 2B-d). SA-induced methylome changes were considerably more frequent than SLT-specific alterations (Fig 2B-e). 3,774 CpGs were hypomethylated upon treatment with SA (alone or in combination with SLT) (Fig. 2B-e), while only 55 CpGs were hypomethylated by SLT (alone or in combination with SA) (Fig 2B-f), showing the considerable potential of SA to induce hypomethylation. Our analysis revealed that SA+SLT modulated CpGs underwent highest hypermethylation in comparison to other treatment conditions (Fig. 2B-d). We also observed proportionately highly abundant hypermethylation events affecting 1,998 CpGs, solely due to SA+SLT exposure (Fig. 2B-d), whereas the hypermethylation events under all other conditions were proportionately far less abundant.

**Figure 2.**
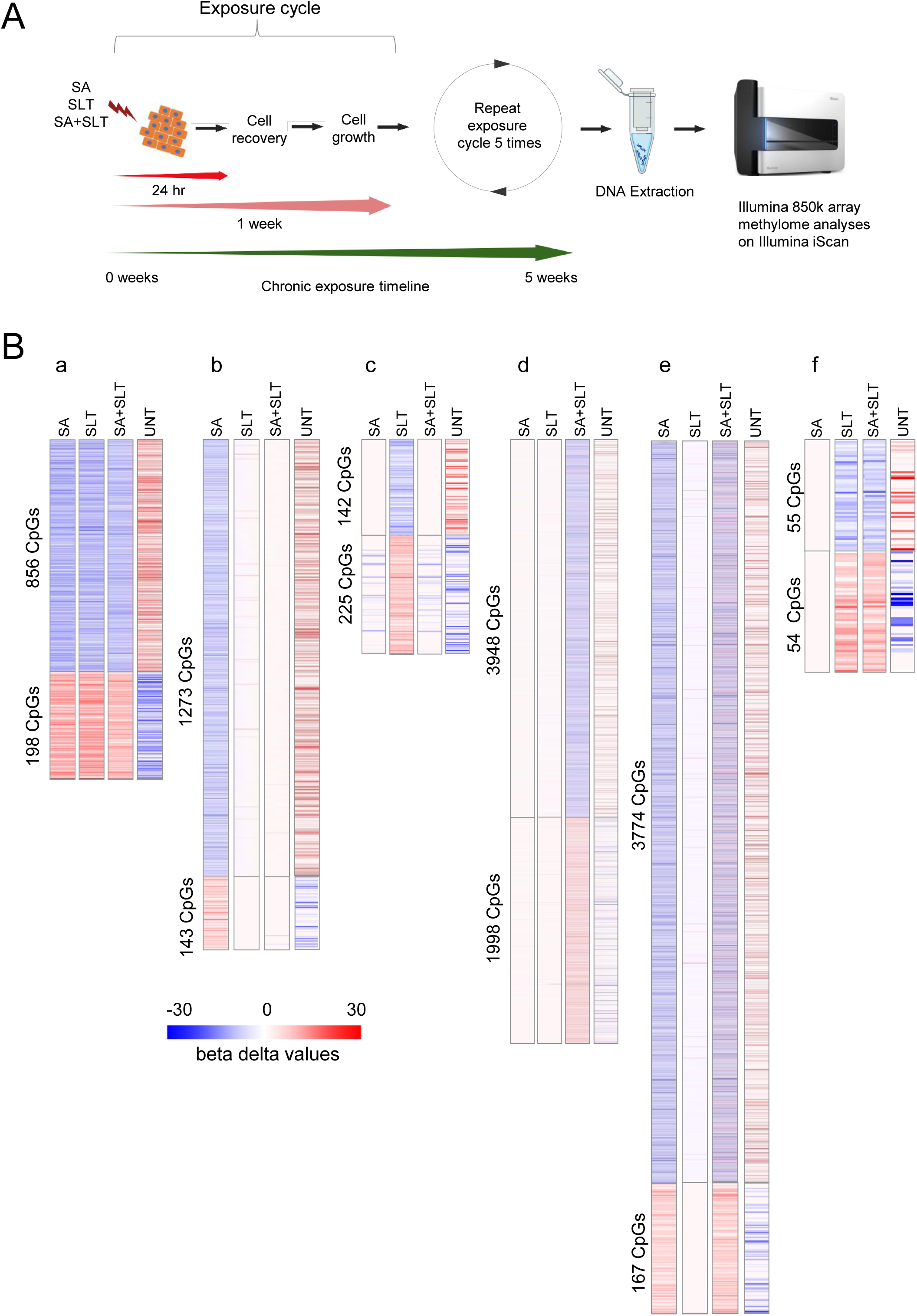
850K Array methylation experiments in NOK cells following chronic exposure: A) Schematic representation of exposure timeline. Some elements of the figure were created using BioRender.com; B) Heat map representation of delta beta values of CpGs under different hypo and hypermethylated conditions: a) SA, SLT, and SA+SLT; b) SA; c. SLT; d) SA+SLT; e) SA and SA+SLT; f) SLT and SA+SLT. Abbreviations: SA - Sodium arsenite (10 µM), SLT - Smokeless tobacco extract (3,000 µg/mL), SA+SLT - Sodium arsenite (7 µM) + Smokeless tobacco extract (1,000 µg/mL), UNT - Untreated time matched control.

Gene ontology analysis based on CpG in promoter regions of neighbouring genes revealed that the extensive hypomethylation induced solely by combined SA+SLT group affected several key biological processes namely, inflammatory response, apoptotic process, cellular response to DNA damage stimulus, chromatin remodelling, DNA damage response, cellular response to reactive oxygen species, among other (Fig 2B-d, Supplementary Table 3d).

SA (alone or in combination) -induced hypomethylation (Fig 2B-e, Supplementary Table 3e) was associated with genes involved in biological processes such as inflammatory response, immune response, apoptotic process, cellular response to DNA damage stimulus, cellular response to reactive oxygen species, and others (Fig 2B-e, Supplementary Table 3e). SLT (alone or in combination)-induced hypomethylation was associated with genes involved in biological processes such as protein ubiquitination and proteasome-mediated ubiquitin-dependent protein catabolic process (Fig 2B-f, Supplementary Table 3f).

In the context of exposure-induced DNA hypermethylation in promoter regions, we observed that hypermethylation caused by SA+SLT co-exposure affected cell-cell signaling, cellular response to tumor necrosis factor, cell division, regulation of cell cycle, and cellular response to DNA damage stimulus, among others (Fig 2B-d, Supplementary Table 4d).

In the case of SA treatment (alone or in combination) and associated hypermethylation (Fig 2B-e, Supplementary Table 4e), we observed affected prominent biological processes including the apoptotic process and cellular response to DNA damage stimulus, among others (Fig 2B-e, Supplementary Table 4e). The prominent biological processes associated with SLT (alone or in combination) induced hypermethylation included proteolysis, protein ubiquitination, and gene silencing by miRNAs, suggesting a gene product turnover/clearance (Fig 2B-f, Supplementary Table 4f).

### 3.3 Arsenic and SLT exposure results in cell cytotoxicity

To investigate the exposure effects on the cell metabolic activity, and to validate the corresponding transcriptomic and DNA methylome remodelling results, we performed MTS assays in NOK and Hupki MEF cells. We observed a gradual decrease in the cell viability due to SA exposure of NOK cells for both tested treatment durations of 24 and 48h, with increasing concentrations (Supplementary Fig. 2A). SLT treatment of NOK cells resulted in a similar dose-dependent reduction in cell viability (Supplementary Fig. 2A). Using the individual cell viability data, the IC_50_ (half-maximal inhibitory concentration) of SA and SLT was established (Supplementary Fig. 3). For both time points, a trend of decreased cell viability with increased exposure concentrations was also observed in the SA + SLT co-exposure group, but the concentrations resulting in a significant drop in viability were much lower as compared to the individual exposure conditions (Supplementary Fig. 2A). We also observed a strong dose dependent cell viability effect in Hupki MEF cells (Supplementary Fig. 2B).

### 3.4 Arsenic and/or SLT exposure affects DNA integrity

The observed decrease in cell metabolic activity due to the tested exposure(s) as well as the results of the transcriptome and DNA methylome remodelling analysis prompted us to investigate whether SA and SLT exposure affects the DNA integrity of the cells, which might in turn might result in decreased cell viability. First, we used γH2AX immunofluorescence to get general understanding of the DNA damage potential of SA and SLT exposure in NOK and Hupki MEF cells. The strongly mutagenic compound aristolochic acid-I (AA) (70) was used as a positive control. The 24h and 48h SA and SLT treatments both individually as well as in combination resulted in γH2AX positivity, like the positive control, while untreated control cells exhibited no to very little γH2AX signal (Fig. 3A). We observed the induction of compromised DNA integrity associated with exposure to the test substances in Hupki MEF cells (Supplementary Fig. 4). DNA damage was not only observed in the context of cytotoxic exposure concentrations, but also when using non-cytotoxic exposure conditions (Supplementary Fig. 4), suggesting that not all detected damage was associated with cytotoxicity.

**Figure 3.**
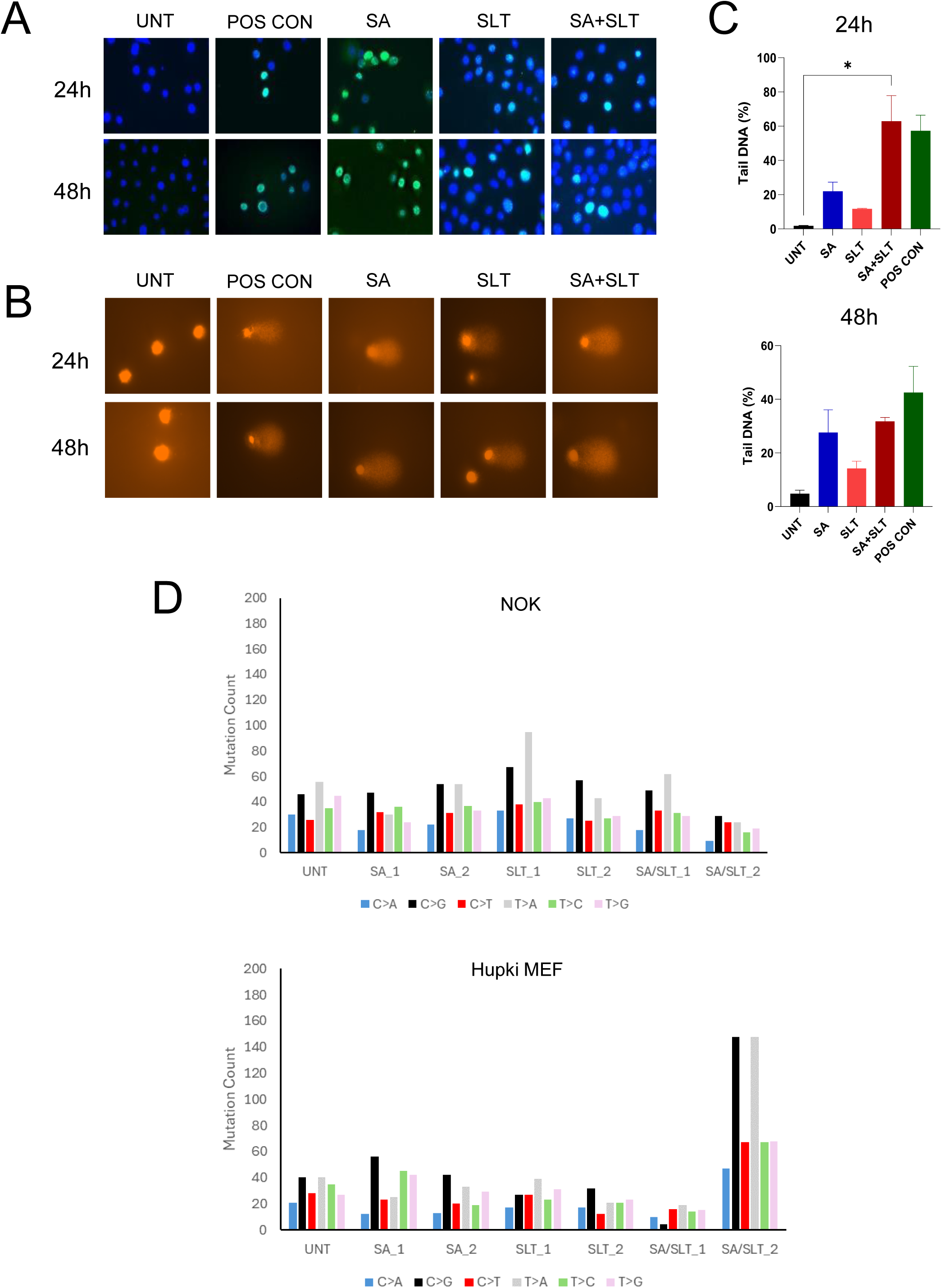
DNA damage response and exome-scale mutation counts: A) γH2AX immunofluorescence (DAPI - blue; Alexa Flour 488 goat anti-rabbit - green) in NOK cells following 24 and 48h exposure. Abbreviations: UNT – Untreated time matched control, , POS CON - Aristolochic acid (100 µM); SA - Sodium arsenite (10 µM), SLT - Smokeless tobacco extract (3,000 µg/mL), SA+SLT - Sodium arsenite (6.5 µM) + Smokeless tobacco extract (750 µg/mL) B) Comet Assay in NOK cells following 24 and 48h exposure: Representative comet images of all the experimental groups. Abbreviations: UNT – Untreated time matched control, POS CON - Aristolochic acid (100 µM), SA - Sodium arsenite (10 µM), SLT - Smokeless tobacco extract (3,000 µg/mL), SA+SLT - Sodium arsenite (10 µM) + Smokeless tobacco extract (3,000 µg/mL); C) Tail DNA percentage after exposure of all the experimental groups in NOK cells. Abbreviations: UNT – Untreated time matched control, SA - Sodium arsenite (10 µM); SLT - Smokeless tobacco extract (3,000 µg/mL), SA+SLT - Sodium arsenite (10 µM) + Smokeless tobacco extract (3,000 µg/mL), POS CON - Aristolochic acid (100 µM); D) Mutation counts in NOK and Hupki MEF cells (For WES exposure dosage information please refer to supplementary tables.). Data (for tail DNA percentage) has been represented as mean ± SE. Statistical analyses (for tail DNA percentage): Kruskal-Wallis test, significance: * =p<0.05. Abbreviations: SA - Sodium arsenite, SLT - Smokeless tobacco extract, AA - Aristolochic acid.

Next, to obtain a more quantitative readout of DNA damage, we performed comet assay in NOK cells (Fig. 3B). Analysis of the comet tails across all the experimental groups revealed a general increase in tail DNA percentage in all treatment groups compared to the time matched controls, with a statistically significant increase detected for the SA+SLT group at the 24h timepoint (Fig. 3C).

The observation of acute DNA damage in the *in vitro* cell models due to SA and SLT (co)exposure prompted us to investigate whether specific mutation spectra may be introduced in chronically exposed Hupki MEF and NOK cells (Supplementary Figure 5, Supplementary Table 7). Whole-exome sequencing analysis of clonally expanded cells did not reveal increases in single base substitution (SBS) numbers or changes in the predominant mutation types in exposed as opposed to untreated control cells (Fig. 3D, Supplementary Table 8, Supplementary Table 9). One of the co-exposed MEF clones exhibited a higher mutation count than the others, possibly due to stochasticity of the senescence bypass process selecting for and resulting in a more mutagenized immortalized clone (59). However, this pattern was not found in any other clones, suggesting that this phenomenon is not generally associated with co-exposure.

### 3.5 Arsenic exposure increases the sub-G1 phase cell population

To further validate several key biological processes and pathways associated with the observed exposure-mediated transcriptome and DNA methylome remodelling (cell cycle regulation, cell cycle checkpoint control, apoptosis, etc.), and to gain additional insight in the acute exposure effects on cell viability and DNA integrity, we performed flow cytometry analysis (Fig. 4). Etoposide was used as a positive control for the analysis, as it induces apoptosis in cells (71). The combined SA+SLT treatment resulted in a significantly increased sub-G1 cell population both following 24h (Fig. 4A, Supplementary Fig. 6, Supplementary Table 10) and 48h of treatment (Fig. 4A, Supplementary Fig. 6, Supplementary Table 11). Although not statistically significant, SA treatment also resulted in an increased sub-G1 cell population after 48h of exposure in comparison to the untreated control (Fig. 4A, Supplementary Fig. 6, Supplementary Table 11). In contrast, SLT treatment did not result in an increased sub-G1 cell population at either exposure timepoint (Supplementary Tables 10 & 11). We did, however, observe significant increase in the G2/M phase cell population due to SLT exposure at 48h (Fig 4A, Supplementary Table11), and also observed a significant increase of G2/M due to SA exposure for 24h (Fig. 4A, Supplementary Table 10).

**Figure 4.**
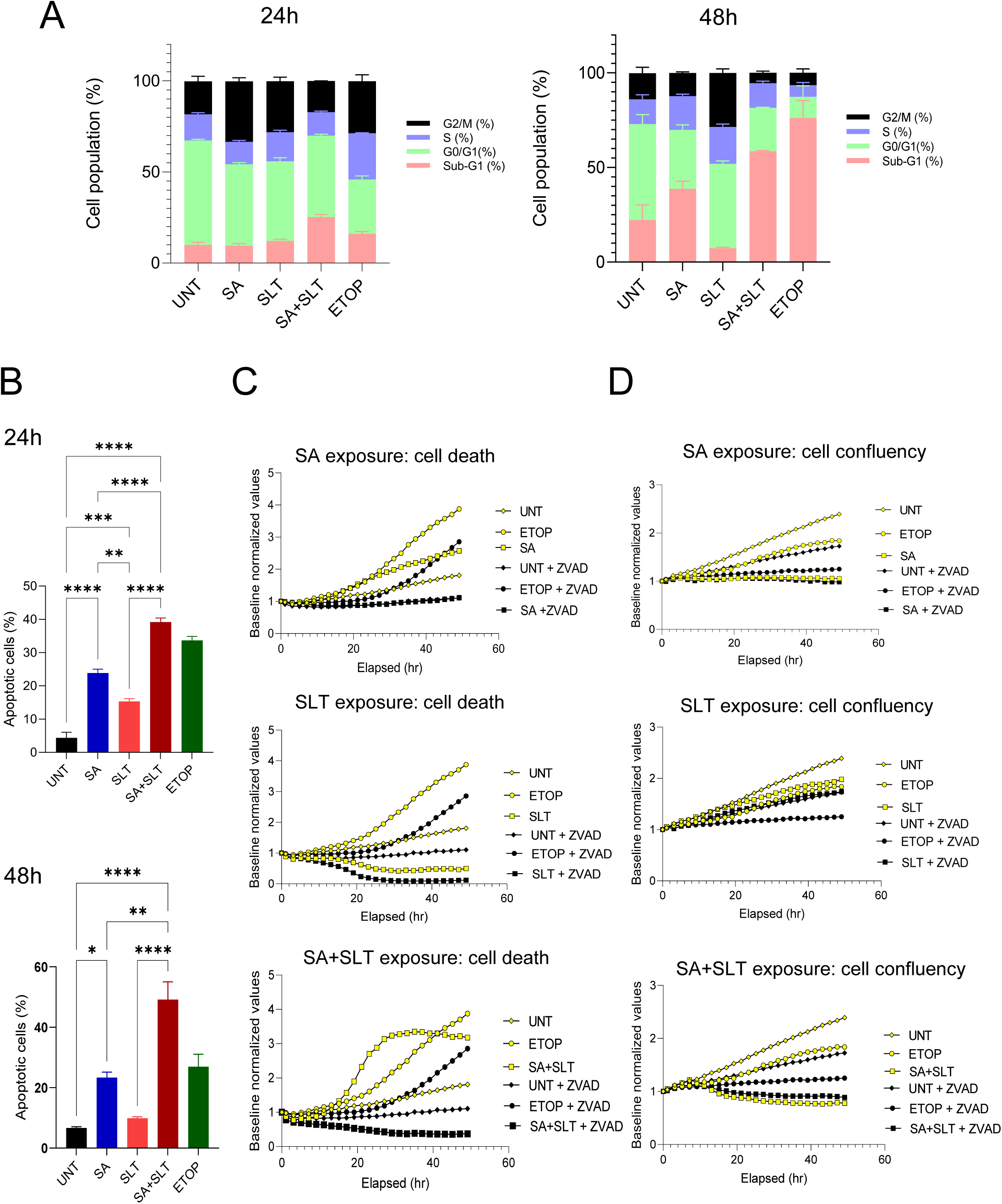
Changes in cellular dynamics in NOK: A) Cell cycle analysis in NOK cells following 24 and 48h exposure; B) Apoptotic cell populations in NOK cells across all the experimental groups following 24 and 48h exposure; C) Cell death monitoring of the experimental groups till 49h by IncuCyte Live-Cell Analysis; D) Cell confluency monitoring of the experimental groups till 49h by IncuCyte Live-Cell Analysis. Abbreviations: UNT - Untreated time matched control, SA - Sodium arsenite (10 µM), SLT - Smokeless tobacco extract (3000 µg/mL), ETOP - Etoposide (100 µM), Z-VAD - Z-VAD-FMK, 1/1000 dilution. Statistical Analyses: One-way ANOVA, Significance: * = p<0.05; ** = p<0.01; *** = p<0.001; **** = p<0.0001

### 3.6 Arsenic exposure increases apoptosis in NOK cells

The high percentage of cell populations with sub-G1 DNA content observed following SA and SA+SLT exposure prompted us to further investigate apoptosis induction using Annexin V staining and flow cytometry. SA exposure induced a significant increase in the percentage of apoptotic cells in comparison to the untreated time matched control group both at 24h and at 48h (Fig. 4B). In comparison to SA, the percentage of apoptotic cells in the SLT treatment group was lower. The percentage of apoptotic cells was significantly higher in the SA+SLT group in comparison to the untreated time matched control group (Fig. 4B, Supplementary Fig. 7).

Next, to confirm the role of apoptosis we exposed NOK cells to the test substances in the absence or the presence of a pan-caspase inhibitor and monitored cell death using Incucyte live-cell analysis. The positive control treatment (etoposide) resulted in a high level of cell death, which was attenuated in the presence of the pan-caspase inhibitor Z-VAD-FMK. Similarly, the amount of cell death induced by SA and SA+SLT treatment was also reduced by Z-VAD-FMK co-administration (Fig. 4C), suggesting that co-exposure of SA and SLT induces cell death via apoptosis.

## 4. DISCUSSION

In this integrative candidate carcinogen exposure study, we employed multi-omic and phenotypic analyses conducted in an experimental model of cultured human normal oral keratinocytes, in order to better understand the mechanisms by which arsenic and smokeless tobacco, both individually and in combination, affect key biological processes that may contribute to cell transformation and oncogenesis.

We observed arsenic exposure resulted in the upregulation of various biological process including apoptosis, response to oxidative stress, and inflammatory response. *MT1E* (which encodes the Metallothionein-1E protein) was found to be one of the genes upregulated due to SA treatment. It is to be noted that the interaction of arsenic with proteins by directly binding to individual cysteine residues, zinc finger motifs, cysteine clusters, etc. plays a critical role in its overall mechanism of toxicity (72). Metallothioneins are a family of cysteine-rich proteins which are confined to the membrane of the Golgi apparatus. They have the capacity to bind various heavy metals like arsenic through the thiol group of its cysteine residues (73) to protect against metal toxicity and oxidative stress, and it has been observed that metallothioneins block the oxidative DNA damage caused by inorganic arsenic exposure (38, 74). *MT1E* expression levels are often found to be altered in cancerous conditions (75, 76).

We observed a significant number of genes upregulated early upon SLT exposure, e.g. *CYP1A1* which belongs to the Cytochrome P450 family of enzymes and is a drug-metabolizing enzyme, is involved in the metabolization of tobacco-related carcinogens (77–80). *CYP1A1* expression levels have been found to be upregulated in cancer cells (81), and gene variants have been associated with tobacco addicted oral cancer cases (82).

Previous studies have shown that arsenic exposure causes changes in DNA methylation in cellular models and in the exposed human populations. (83, 84). Studies have revealed that arsenic-mediated widespread DNA hypomethylation is associated with promoter activation and carcinogenesis (85–87). Our observation of significant hypomethylation induced by SA or SA/SLT co-exposure and the deregulation of common biological processes and pathways on the methylome and transcriptome level due to SA and SLT effects points to an integrated exposure-mediated molecular reprogramming that contributes to specific cellular phenotypes with potential roles in carcinogenesis.

We observed considerable overlap of biological processes and pathways associated with transcriptomic and DNA methylome changes in response to SA exposure (Supplementary Table 5, Supplementary Table 6). Some of the commonly deregulated processes included the immune system, inflammatory response, chromatin, apoptotic process, pathways in cancer, cellular response to DNA damage stimulus, chromatin organization, DNA Repair, response to oxidative stress, regulation of TP53 activity, cell cycle checkpoints and others (Supplementary Table 5).

The pronounced cytotoxicity and DNA damage, observed especially upon arsenic and SLT co-exposure, are supported by recent studies that demonstrated the cytotoxic (88–90) and genotoxic (91–93) potential of arsenic treatment in cells. Similarly, cytotoxicity of various types of SLT in *in vitro* test models has been documented (94, 95). Studies in exposed experimental animal test models and evaluation of samples from exposed human populations have confirmed the DNA damage causing potential of arsenic and SLT co-exposure (50, 17, 51, 18, 52, 96). In line with the upregulation of oxidative stress response programs in our multi-omics analysis, chronic exposure to arsenic and SLT in experimental animal *in vivo* models results in severe induction of oxidative stress (50, 51). The overpowering of DNA repair and antioxidant mechanisms by reactive oxygen species can result in oxidative stress and damage, which might play an overall critical role in chemical carcinogenesis (97).

We observed higher sub-G1 population due to arsenic and combined exposure at the 48h changes signifying increase in cell death due to co-exposure. Further, Annexin V PE assay analysis in the cells revealed significant increase of apoptotic cells in the arsenic and the combined groups following acute exposure. Previous studies have reported the induction of apoptosis or autophagy due to arsenic exposure (98). We also demonstrated that Z-VAD-FMK was able to rescue cells from exposure mediated cell death. Altogether, the apoptosis induced by the SA or SA+SLT exposure, as validated by multiple assays used, is a likely cause of the observed cytotoxicity and DNA damage.

The observed short-term genotoxicity did not translate to long-term, exome-scale mutagenic effects under any (co)exposure condition. Based on the analysis of thousands of cancer genomes and exomes, a characteristic SBS mutational signature (SBS29), found predominantly in oral squamous cell carcinoma and hepatocarcinoma, has been associated with tobacco chewing (99). It is to be noted that the *sadagura* form of SLT contains various active chemical compounds (18), some of which e.g., estragole, when metabolized, can form DNA adducts, which may result in mutagenicity if they remain unrepaired (100). The absence of an increase in SBS counts in the current study suggests that the compounds in the SLT extract do not induce such mutagenesis, potentially due to insufficient metabolization into active compounds or due to DNA adduct repair by NOK or Hupki MEF cells. Our analysis focused on SBS mutations, however other mutation types such as small insertions and deletions, copy number variations and structural variations can contribute to carcinogenesis. For example, arsenic-induced skin lesions exhibit genome wide copy number variation (101). Moreover, various types of genetic alterations such as indels have been linked to tobacco and betel quid use-associated oral cancer (102). Considering these observations, we cannot rule out the contribution of other mutational mechanisms to SA and SLT-mediated genetic alterations.

The global molecular changes observed in our short- and long-term exposure settings suggest the common induction of gene expression programs associated with cell death and inflammatory responses upon arsenic and SLT (co)exposure (Fig. 5). The identification of molecular programs associated with cell death is in keeping with the observed phenotypes and this might explain the observed DNA damage. The presented catalogue of molecular programs and events reported here may have implications for monitoring of populations in South Asia exposed to high levels of arsenic via groundwater while habitually using one of various forms of SLT preparations. The extensive deregulation of molecular and phenotypic programs following exposure to SA and/or SLT warrant future systematic design of molecular epidemiology studies in humans, as well as generating evidence base for reducing the exposure to arsenic and smokeless tobacco products and consequently lessening the risk of oral cancer formation.

**Figure 5.**
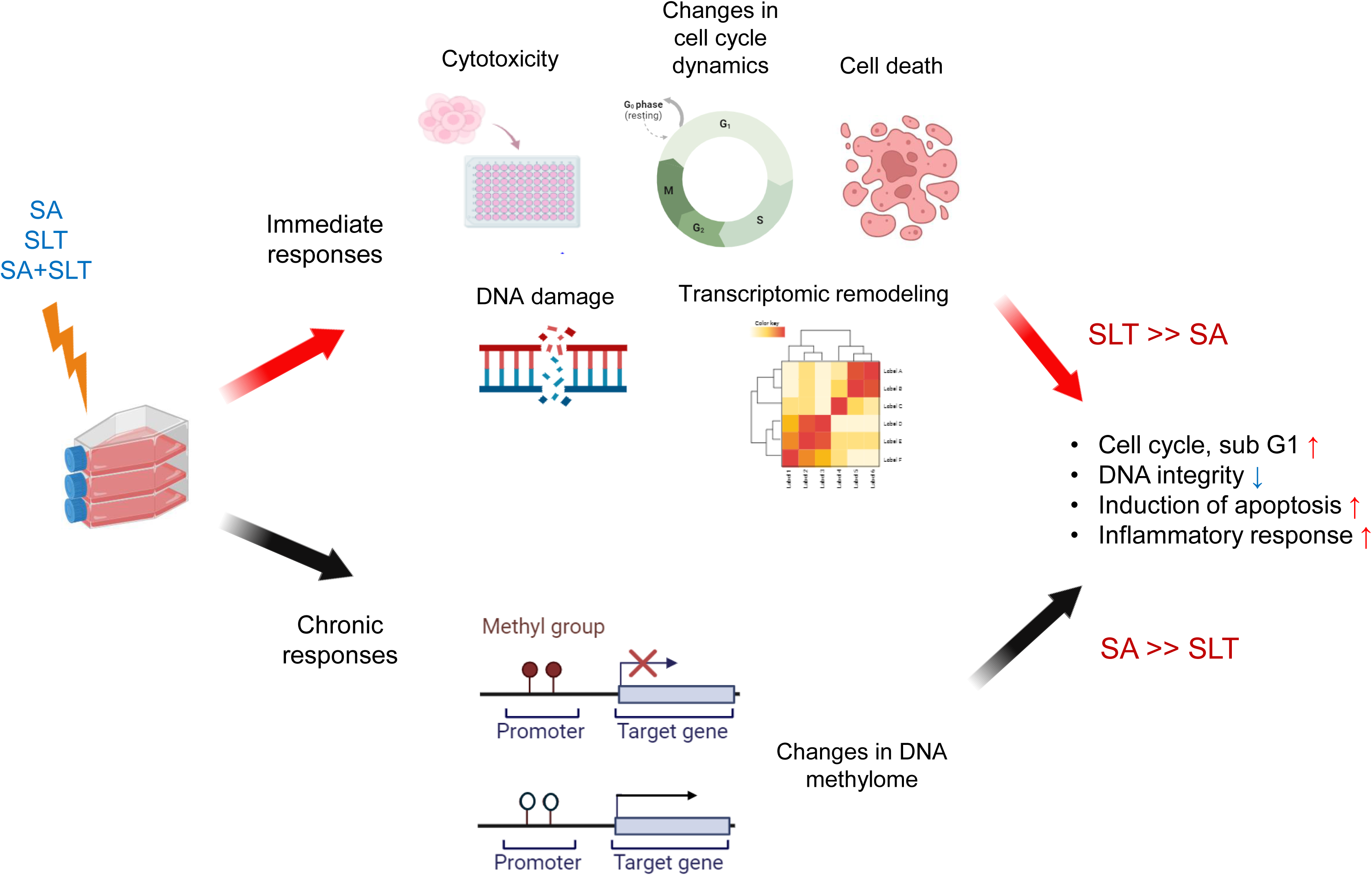
Based on our in-vitro findings, possible mechanism(s) by which potentially arsenic and smokeless tobacco co-exposure can cause induction of carcinogenesis. The figure was created using elements from BioRender.com.

## DISCLAIMER

Where authors are identified as personnel of the International Agency for Research on Cancer / World Health Organization, the authors alone are responsible for the views expressed in this article and they do not necessarily represent the decisions, policy or views of the International Agency for Research on Cancer / World Health Organization.

## ACKNOWLEDGMENTS

The work reported in this paper was undertaken during the tenure of an IARC Postdoctoral Fellowship (to Samrat Das) at the International Agency for Research on Cancer, Lyon, France. The Hupki MEF exposure system had been established with the support from INCa-INSERM (Plan Cancer 2015) and NIH/NIEHS (1R03ES025023-01A1).

## DATA AVAILABILITY STATEMENT

The original methylome array and RNA sequencing data are publicly available from the NCBI’s Gene Expression Omnibus (GEO), under the respective accession numbers GSE269103 and GSE269688.

---

**Supplementary Figure 1.**
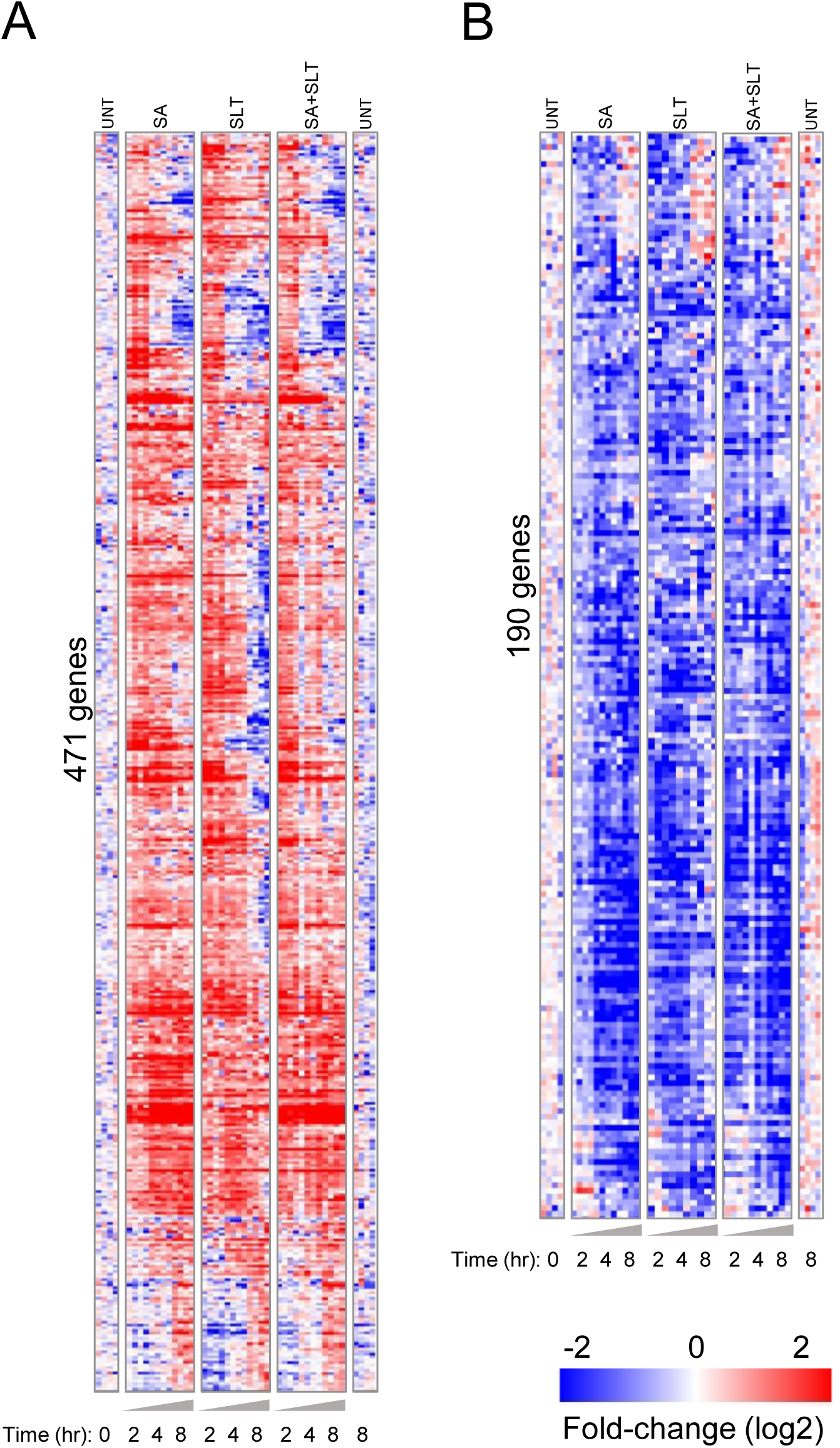
General gene regulation patterns irrespective of treatment conditions: A) General response, upregulated genes; B) General response, downregulated genes. Abbreviations: UNT -Untreated time matched control, SA - Sodium arsenite (10 µM), SLT - Smokeless tobacco extract (3000 µg/mL). SA+SLT - Sodium arsenite (6.5 µM) + Smokeless tobacco extract (750 µg/mL).

**Supplementary Figure 2.**
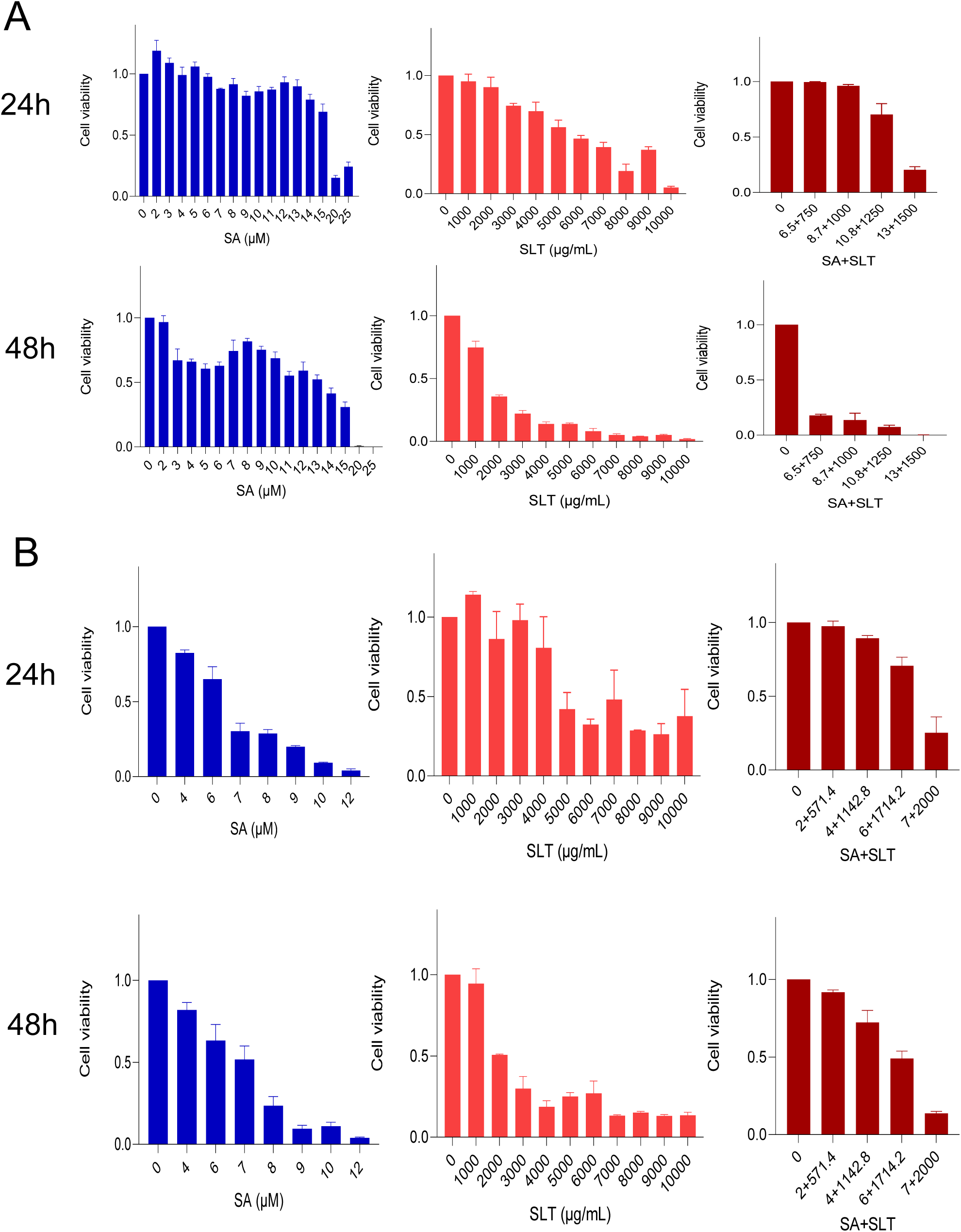
Cytotoxic response in: A. NOK and B. Hupki-MEF cells. Abbreviations: SA - Sodium arsenite, SLT - Smokeless tobacco extract, SA+SLT - Sodium arsenite + Smokeless tobacco extract.

**Supplementary Figure 3.**
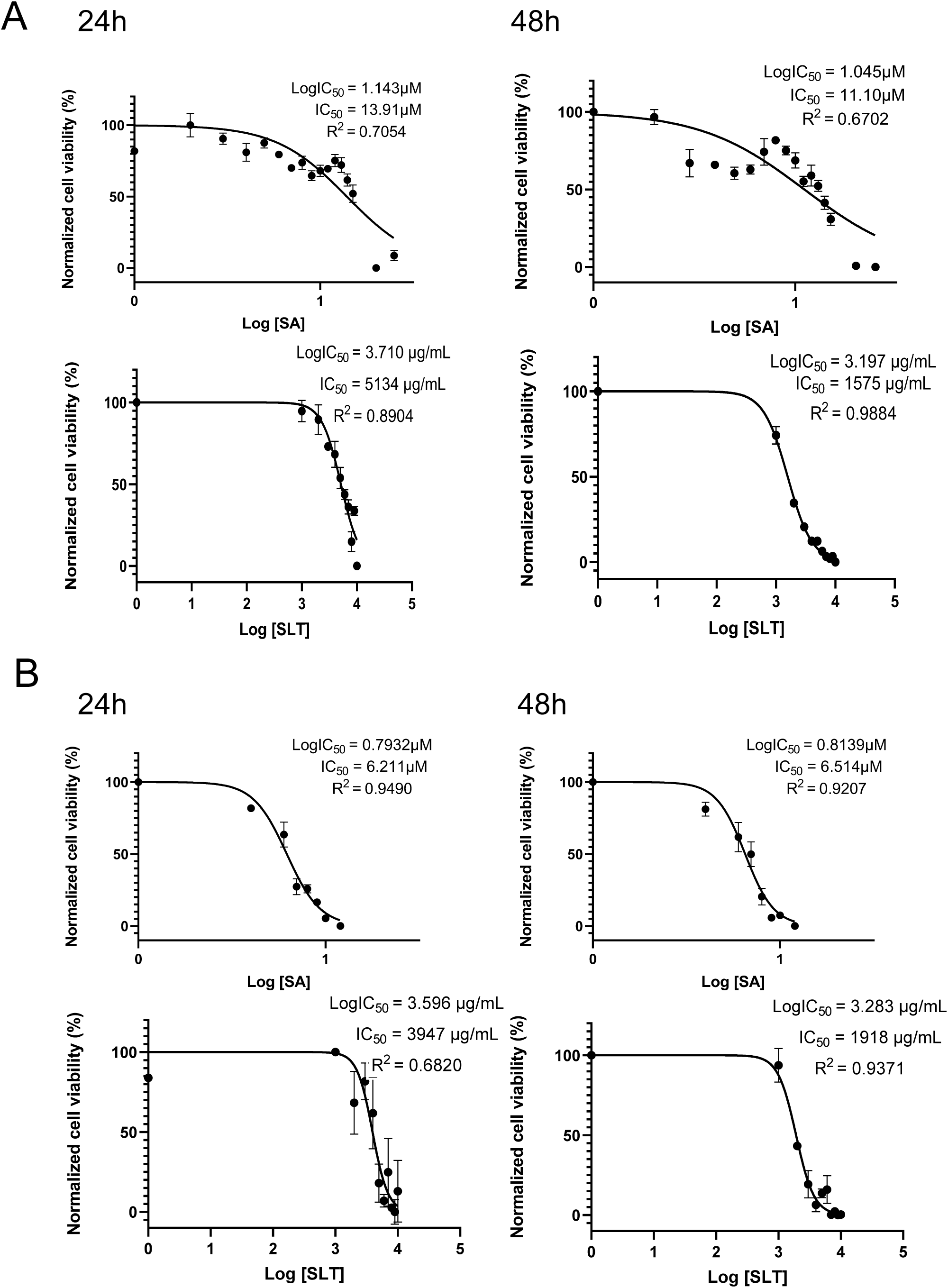
Half maximal inhibitory concentrations, IC_50_ analysis in: A) NOK and B) Hupki-MEF cells. Abbreviations: SA - Sodium arsenite, SLT - Smokeless tobacco extract. Log IC_50_ : logarithmic half maximal inhibitory concentrations. R^2^ : measure of goodness of fit.

**Supplementary Figure 4.**
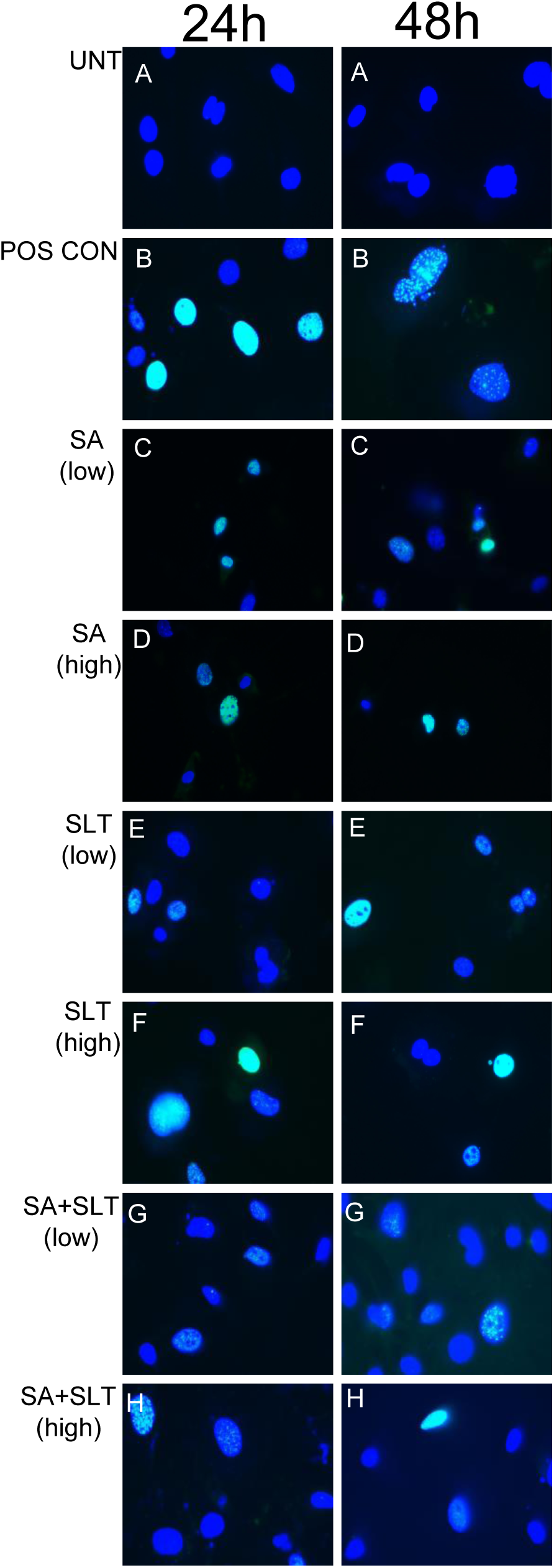
Genotoxic response (Gamma H2AX immunofluorescence) in Hupki-MEF cells. A. UNT, Untreated time matched control, B. POS CON, Positive control, AA 100µM, C. SA (low), SA exposure (4µM), D. SA (high), SA exposure (7µM), E. SLT (low), SLT exposure (1000 µg/mL), F. SLT (high), SLT exposure (4500 µg/mL), G. SA+SLT (low), SA+SLT exposure (2µM+571.4 µg/mL), H. SA+SLT (high), SA+SLT exposure (6µM+1714.2 µg/mL).

**Supplementary Figure 5.**
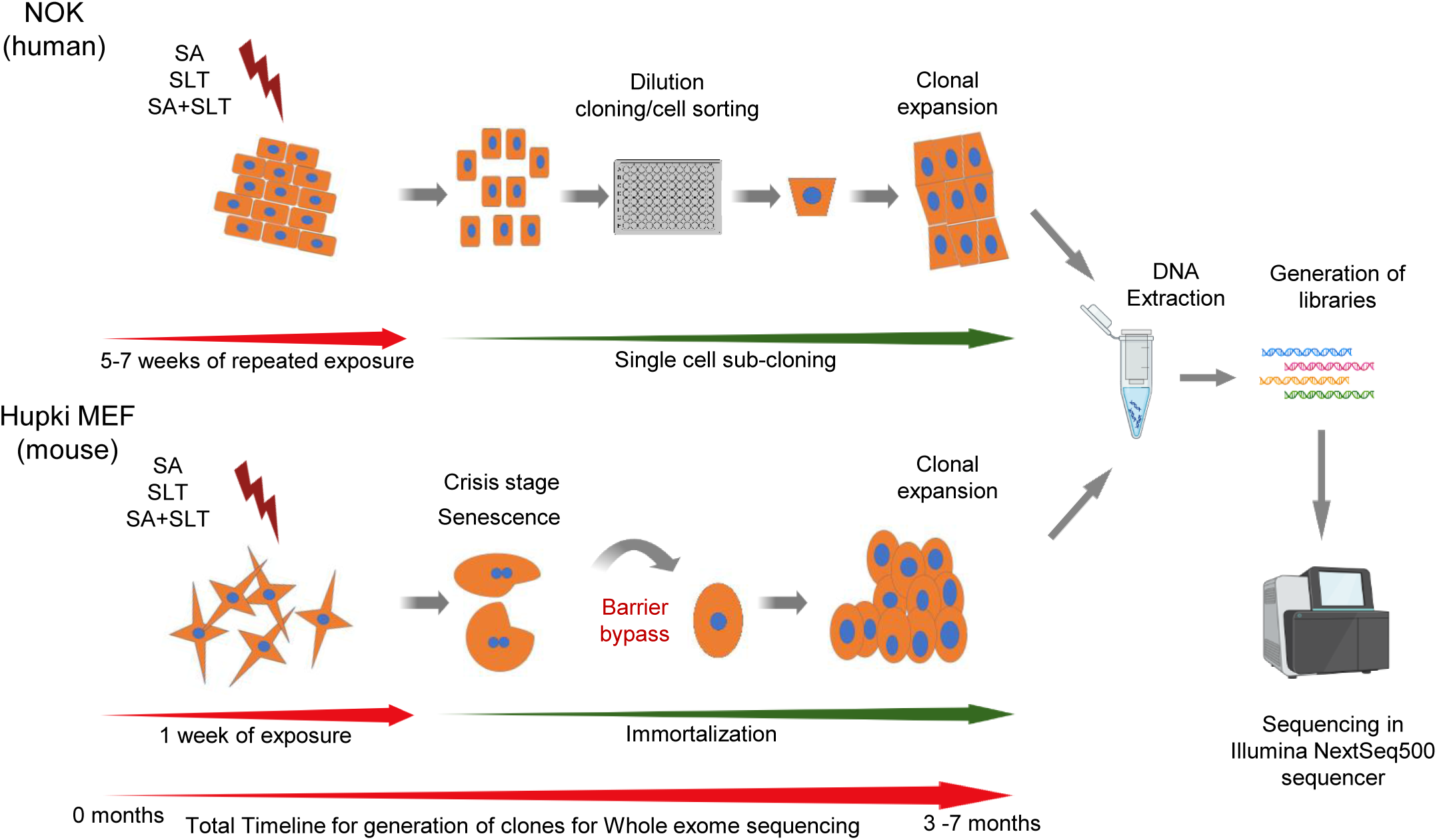
Schematic representation of the chronic exposure and generation of clones for whole exome sequencing in cells. Some elements of the figure were created using BioRender.com.

**Supplementary Figure 6.**
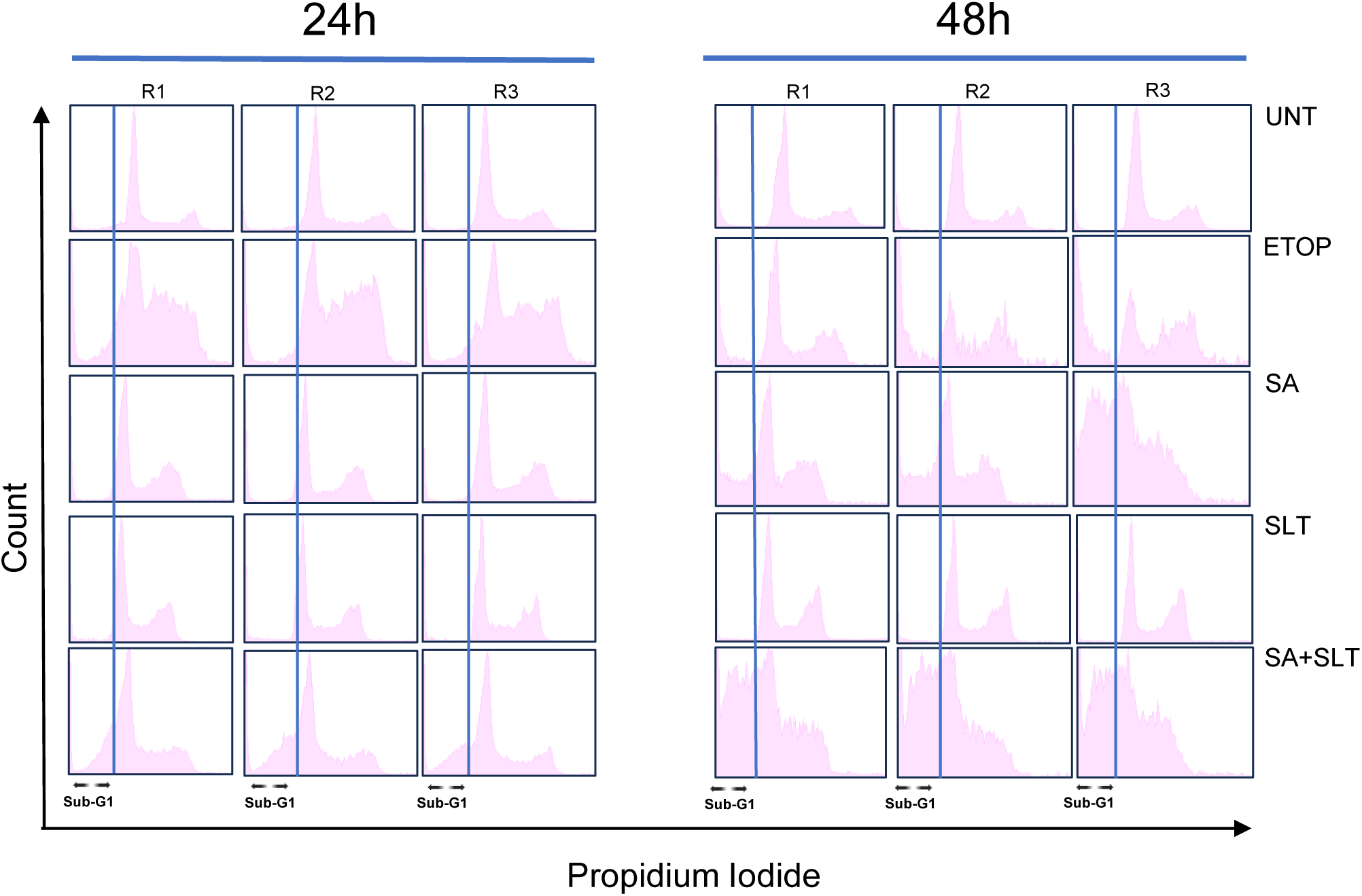
Cell cycle analysis by flowcytometry in NOK cells: Biological replicate PI histogram plots of all the experimental groups (Experiment was performed in triplicates per experimental group). Abbreviations: UNT - Untreated time matched control, ETOP – Etoposide (100µM), SA - Sodium arsenite (10 µM), SLT - Smokeless tobacco extract (3000 µg/mL), SA+SLT - Sodium arsenite + Smokeless tobacco extract. R - Replicates.

**Supplementary Figure 7:**
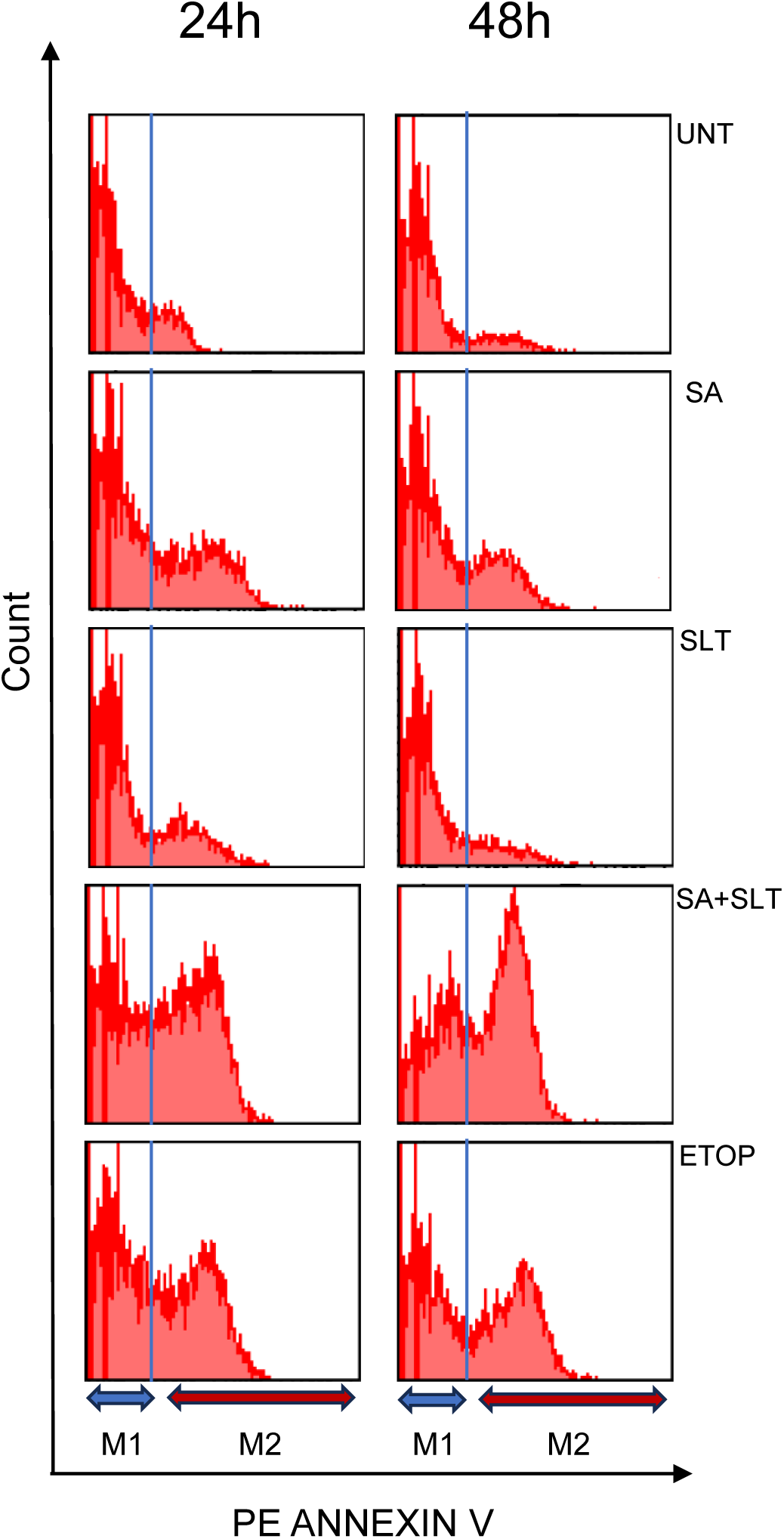
Apoptosis analysis in NOK cells: Representative Annexin V PE versus count histogram plot output images. Abbreviations: UNT – Untreated time matched control, SA - Sodium arsenite (10 µM), SLT - Smokeless tobacco extract (3000 µg/mL), SA+SLT - Sodium arsenite + Smokeless tobacco extract, ETOP – Etoposide (100 µM), M1 - Cells that were viable and not undergoing apoptosis, M2 – Cells that are undergoing apoptosis.

